# Mechanisms of long term non-reinforced preference change: functional connectivity changes in a longitudinal functional MRI study

**DOI:** 10.1101/2025.04.08.647732

**Authors:** Alon Itzkovitch, Shiran Oren, Sidhant Chopra, Alex Fornito, Tom Schonberg

## Abstract

Behavioral change studies mostly focus on external reinforcements to modify preferences. Cue-approach training (CAT) is a paradigm that influences preferences by the mere association of stimuli, sensory cues, and a rapid motor response, without external reinforcements. The behavioral effect has been shown to last for months after less than one hour of training. Here, we used a modified version of CAT by changing the neutral-cue to a number that represented a monetary amount of reward that the participants accumulate (i.e. incentive-cue). After a single training session, we compared behavioral performance and functional connectivity (FC), as measured using functional magnetic resonance imaging, between two groups, one receiving a neutral-cue and the other receiving an incentive-cue, at 5 time points across one year. We replicated the maintenance of behavioral changes after 6-months for the non-reinforced neutral-cue participants, but not for the reinforced group. The reinforced training group showed higher FC within the limbic system, whereas the non-externally reinforced group showed higher functional connectivity within and between default-mode and dorsal-attention networks. Our findings offer putative neural correlates for both reinforced and non-reinforced preference changes that are maintained over time and which could be implemented in future behavioral change interventions.

**Significance Statement:** The current work examines the neural mechanisms of non-externally reinforced preference change and its maintenance over time, using both a neutral cue and a modified version of the cue-approach training paradigm. While both groups initially exhibited preference shifts, only the non-reinforced group maintained preference changes over one year, suggesting an enduring internal reinforcement mechanism. We identified distinct patterns in functional connectivity related to behavioral maintenance, find that non-external reinforcement increases the connectivity of the default mode network while external reinforcement elevates connectivity between limbic areas. These findings enhance our understanding of sustainable behavior change and advocate for non-external reinforcement in behavioral interventions.

## Introduction

Behavioral change is essential to addressing public health challenges and interventions, often achieved through training paradigms that shift preferences and encourage desirable behaviours (Michie et al., 2011). A century ago, Pavlov (Pavlov & Anrep, 1927) showed that pairing a sensory cue with external reinforcement could elicit the same response to the cue alone after training. The Pavlovian conditioning concept laid the foundation for learning theory and a myriad of external-reinforcement behavioral paradigms, where rewards like monetary prizes drive behavior change (Dayan and Niv 2008). Nonetheless, inducing long-term changes in behaviour has proven challenging for the field, with extant research indicating that reinforcement-based strategies often results in transient effects, with behaviors diminishing once rewards are no longer provided (Bouton, 2002; Christiansen et al., 2007; Prochaska et al., 2004).

In recent years several studies have examined non-externally reinforced preference change paradigms. For example inhibitory-responses (Jones et al., 2016) and Go-NoGo (Veling, Lawrence, et al., 2017) tasks, where participants repeatedly withhold responses to specific stimuli, have been shown to modify subjective preferences to trained items without external reinforcements. More recently, the cue-approach training (CAT) paradigm (Schonberg et al., 2014) has been used to show that the mere association between a speeded motor button-press response and a sensory cue, coupled with a visual stimulus, leads to preference modification towards trained items, without external reinforcements. Multiple studies have replicated this behavioral effect (Aridan et al., 2019; Bakkour et al., 2016, 2017, 2018; Botvinik-Nezer et al., 2020, 2021; Chen et al., 2016; Salomon et al., 2020, 2022; Veling, Chen, et al., 2017; Zoltak et al., 2018), and demonstrated its generalization across various stimuli and cue types (Salomon et al., 2018). Moreover, the effect of one-hour CAT training can lead to preference changes lasting up to 6-months (Salomon et al., 2018).

Several mechanisms have been offered to explain the behavioral effects of the CAT (Bakkour et al., 2017; Botvinik-Nezer et al., 2020; Salomon et al., 2020; Schonberg et al., 2014). Botvinik-Netzer et al. (2020) suggested it is driven by a decrease of top-down attentional processes coupled with an enhancement of bottom-up processes driven by training on the task (Botvinik-Nezer et al., 2020). A recent meta-analysis using a Bayesian computational learning model showed a positive correlation between the shift in reaction-time during training and the subsequent behavioral change effect. The mechanism offered for preference change was linked to internal reinforcements, whereby the cue transformed from a mere signal into a reinforcer (Salomon et al., 2022).

The observed long-lasting behavioral changes of the CAT (Salomon et al., 2018), which is not seen with most reinforcement-based strategies, suggest the involvement of distinct mechanisms supporting preference modification. Schonberg and Katz (2020) offered the existence of two distinct neural circuits for reinforced and non-reinforced behavioral changes. They proposed that a dopamine dependent ventral-value pathway, involving amygdala and hippocampus, underlies externally reinforced preference change, while a dorsal-value pathway, involving occipital and parietal regions, underlies non-externally reinforced preference changes. Both circuits implicate striatum and pre-frontal regions, due to the known role they play in reward and value processing (Delgado, 2007; Kringelbach, 2005; Rangel et al., 2008).

To date, no study has directly compared reinforced and non-externally reinforced paradigms, nor their underlying mechanisms. Here, we aimed to bridge this gap by modifying the unique CAT task and replacing the non-incentivized sensory cue with a monetary cue to create an incentive-cue group. We used functional magnetic resonance imaging to test differences between incentivized vs. non-incentivized groups in behavioral change and inter-regional functional connectivity immediately following one hour of training and during four subsequent follow-up timepoints over the following year (i.e., 1,3,9, and 12 months) to assess the persistence of preference modifications. We hypothesized and preregistered that a similar behavioral effect would be observed for both groups immediately after training, with higher decay rates for the incentive-cue group due to the removal of the reinforcer. No specific hypotheses were pre-registered as to the neural systems involved.

## Materials and Methods

### Participants

We collected data from 107 valid participants at the first time point (56 in the neutral group, 51 in the incentive-cue group); at the second time point, participant numbers were 40 (neutral) and 36 (incentive-cue); and at the final follow-up session, numbers were 15 (neutral) and 18 (incentive-cue).

The original sample size for this study was based on a power analysis of 80% power with an alpha of 0.05 that was calculated on previously collected data in our lab (Botvinik-Nezer et al., 2020), which examined responses to high-value Go items in the vmPFC. We planned to scan sixty-two participants in each group (124 overall). Dropout was predicted to be 25% over the year; therefore, we expected ∼45 participants in each group for the final follow-up session. However, due to the COVID-19 pandemic our final sample size was smaller. Demographic details are shown in Table 1.

**Table 1.** Demographic details. For each group at each timepoint, the table describes: 1) the number of participants (and number of females within the sample size); 2) the mean number of days between the current session and the first one (standard deviation in parentheses); 3) the mean age of participants (standard deviation in parentheses).

| Timepoint | Neutral-Cue group |  |  | Incentive-Cue group |  |  |
| --- | --- | --- | --- | --- | --- | --- |
|  | Participants<br>(Females) | Days after 1<sup>st</sup><br>session (std.) | Age (std.) | Participants<br>(Females) | Day after 1<sup>st</sup><br>session (std.) | Age (std.) |
| **1** | 56 (29) | 0 (0) | 26.1 (4.8) | 51 (19) | 0 (0) | 26.7 (4.3) |
| **2** | 40 (23) | 44.5 (23) | 26.1 (5.3) | 36 (15) | 46 (27.8) | 27.3 (5) |
| **3** | 31 (19) | 102.4 (26) | 25.7 (4.5) | 26 (10) | 106.6 (22.5) | 27.9 (4.8) |
| **4** | 21 (14) | 306.3 (36.5) | 26.8 (5.4) | 22 (9) | 296 (28.3) | 28.1 (5.4) |
| **5** | 15 (12) | 391 (27.3) | 24.8 (4.6) | 18 (6) | 377.9 (16) | 28.1 (5.1) |

### Study Design

#### Stimuli

The stimuli used in the CAT task were created in our laboratory and composed of 80 popular Israeli snack food items, presented on a black background. All snacks were available at main stores in Israel and cost no more than $2.7 USD (equal to 10 NIS). Stimuli were presented using Matlab and Psychtoolbox-3 (Pelli, 1997). The stimuli are a subset of bigger database that can be found at https://www.schonberglab.sites.tau.ac.il/resources.

#### Image Acquisition

Participants were scanned using a Siemens Prisma 3T magnetic resonance imaging (MRI) scanner at Alfredo Federico Strauss Center for computational neuroimaging at Aviv University. Structural images were acquired using MPRAGE (Brant-Zawadzki et al., 1992) and FLAIR (Bakshi et al., 2001) protocols. A multiband EPI (echo-planar imaging) sequence (multi-band acceleration factor = 4; TE = 30.0 ms; TR = 1,200 ms; flip angle = 72°; voxel resolution = 2 mm^3^) was used for the task-fMRI. For field mapping, a spin-echo protocol was used. For resting-state scans, we used an EPI protocol (TE = 30.4 ms; TR = 750 ms; flip angle = 52°; voxel resolution = 2*2*2 mm), for 350 volumes in both AP and PA directions (full runs in both directions).

#### Procedure

Cue-approach training (CAT) is a paradigm for non-external reinforced behavioral change first introduced in 2014 (Schonberg et al., 2014). The task is composed of three phases: (1) an assessment of individuals’ initial willingness to pay for items; (2) a training phase; and (3) a probe phase, in which participants are asked to make binary choices between trained and untrained items that have similar initial willingness to pay scores. Full details are provided below (see *Cue approach task section*).

During the task, the mere association between a rapid motor response to visual stimuli paired with a cue leads to preference modification (Aridan et al., 2019; Bakkour et al., 2016, 2017, 2018; Botvinik-Nezer et al., 2020, 2021; Salomon et al., 2018, 2020; Schonberg et al., 2014). The task was initially performed with an auditory cue and snack food item stimuli (Schonberg et al., 2014). In the current study, we performed the CAT procedure with visual cues while participants were scanned with fMRI. The experiment included five imaging meetings (sessions) over one year: follow-up sessions (session 2-5) were planned to be conducted after 1,3,9 and 12 months from the first meeting. See Figure 1 for a general layout of the task.

**Figure 1.**
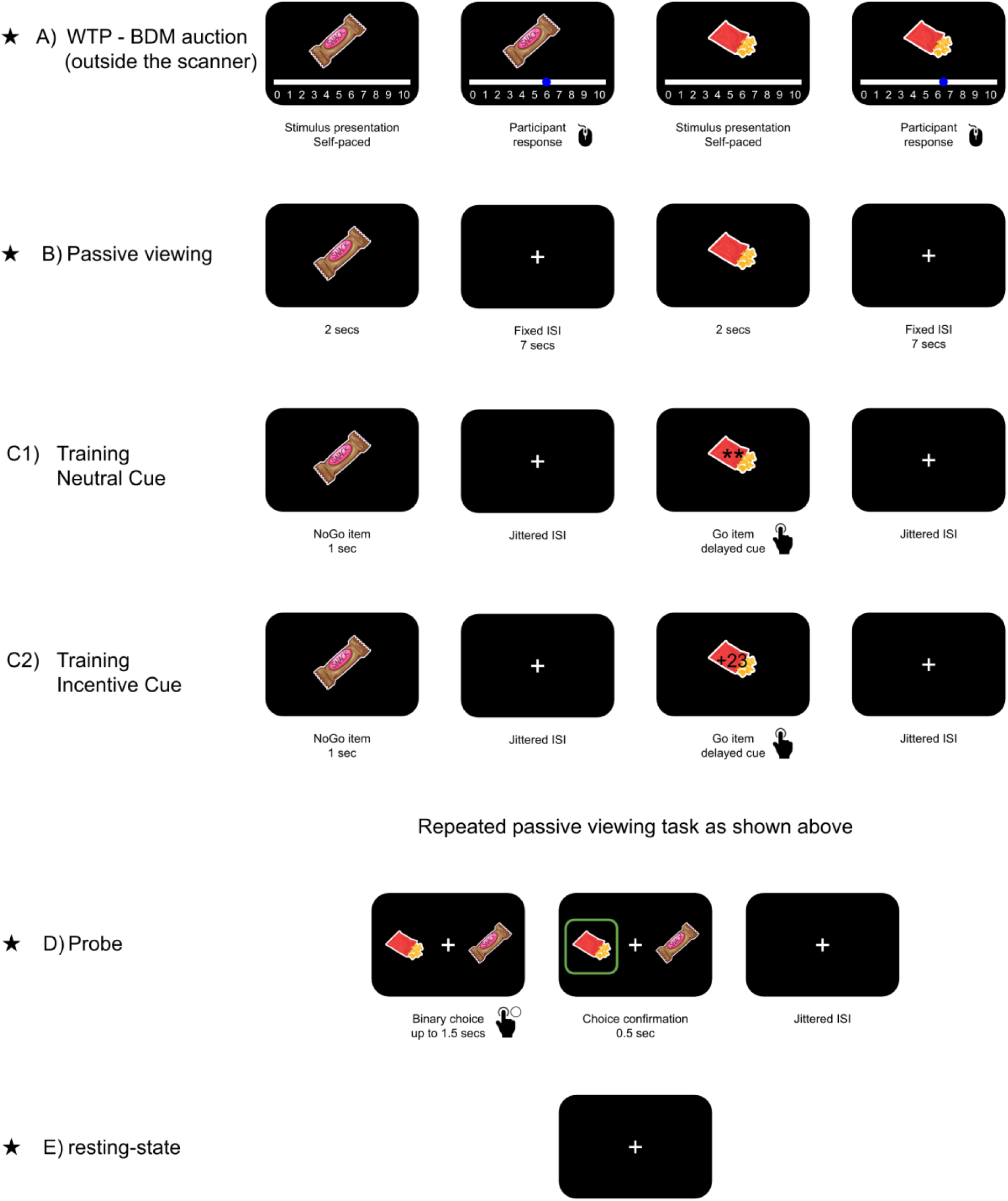
sequence of events. The figure demonstrates the experiment during the first session. *(A)* a Becker-DeGroot-Marshcak (Becker et al., 1964) auction task (outside the scanner): participants willingness to pay (WTP) for each of the snacks was measures. (B) Passive viewing: snacks were shown one by one. (C1) Training task for the neutral-cue group: Items were shown one by one on the screen. Some of the items were associated with a cue (Go items). The cue was two asterisks presented ∼750ms after the stimuli appeared. Participants were asked to press a button as fast as they could in response to the cue. (C2): Training task for Incentive -cue: Like C1, except the cue was a number that represented the amount of money in which participants could win at the end of the experiment. (D) Probe: binary choice between Go items and NoGo items with the same initial value. (E) resting-state scan. The follow-up phases are marked with an asterisk. Figures adapted from Botvinik-Nezer et al. 2020. Snacks icons made by <u>Freepik</u> from www.flaticon.com.

Participants were randomly assigned to two groups: the neutral-cue group (NC) and the incentive-cue group (IC). Both groups performed an identical procedure. The only difference was the visual cue presented during the training phase during the training session (see figure 1C). In follow-up sessions (i.e., at 1, 3, 9, and 12 months), participants performed only the probe task to assess the sustainability of training effect, followed by a memory task.

#### Cue approach task (CAT)

Figure 1 presents an overview of the entire experimental design. The procedure was based on previous CAT studies, with a visual instead of auditory cue (Botvinik-Nezer et al., 2020; Salomon et al., 2020; Schonberg et al., 2014).

### Session 1

1. ***Willingness to pay:* (15 minutes; outside the scanner.):** In order to obtain the participant’s subjective initial willingness to pay (WTP) preferences for 60 snack food items, we used the Becker-DeGroot-Marschak procedure (BDM, (Becker et al., 1964)). Participants were asked to fast for four hours prior to arriving to the laboratory. First, participants were given 10 NIS and asked to participate in an auction. Each snack food item was presented and rated on an analogue scale from 1 to 10 using the mouse, describing the subjective willingness to pay to a given item, without a time limit. Then, the sorted ratings were used to determine each persons’ snack ranking, reflecting their preferences.
2. ***Passive viewing task* (10 minutes; inside the scanner):** This task aimed to study neural activation while participants passively viewed the items without any additional manipulations (Botvinik-Nezer et al., 2019). A total of 40 snacks––20 high-value and 20 low-value items based on individualized preferences––were chosen based on the WTP phase according to each person’s own subjective ratings. Items from the preference ranking list of each participant ranked 3-22 were considered as high-value items and items ranked 39-58 were considered as low-level items. Those items were also presented during the training phase (see section 3 below). To ensure that participants were attending to the items they were instructed to look at the items and count how many were one or many (i.e. a “Mars” bar is single, whereas Doritos are many). On each trial, a random item was presented from the subset for 2 seconds, followed by a fixed interval of 7 seconds. Two runs were conducted: each run presented all 40 items (each item was presented twice in the task, 80 trials over 2 runs). Each run started with a 2 second fixation and ended with 7seconds post-run fixation completion to the last run fixation.
3. ***Training task* (30 minutes; inside the scanner):** the same 40 items from the passive viewing phase were presented one by one for 1 second, followed by a jitter interval of 3.5 seconds, on average (SD = 1.21 seconds, range of 1 - 7 seconds, 1 second interval). Each run started with 2 seconds of a pre-run fixation and ended with 6 seconds post-run fixation completion to the last run fixation. Thirty percent of the items were coupled with a visual cue, and termed Go items. The remaining items were not associated with a cue and were termed NoGo items. Participants were instructed to press a button as fast as possible when a cue appeared, before the stimulus disappeared. Twelve runs were conducted, with random order of the stimuli at each run, resulting in 480 trials in total. For both NC and IC groups, there was no external indication whether the press was performed on time. The timing of the cue appearance after a stimulus-presentation was calculated using a stepwise procedure to obtain 75% successful button presses. Four subsets of stimuli were used in this phase, two subsets with the same mean value of high-value items (ranked 3-22 in the WTP phase) and two subsets of low value items (ranked 39-58 in the WTP phase). One high-value and one low-value subset were associated with a cue (Go items).
4. *Repeated passive viewing task* (10 minutes; inside the scanner): The same task as section 2 above was repeated.
5. ***Probe task* (15 minutes; inside the scanner):** In this phase, on each trial two similarly initial valued items were pitted against each other. One of the items was previously a Go item (during training) and the other was a NoGo item. The items were located randomly on both sides of the screen. A total of 144 comparisons were presented during two separate runs. On each run, 36 high-value trials and 36 low-value trials were conducted. Also, four sanity checks were added to each run, comparing high- and low-value NoGo items (to test if participants were consistent with the BDM initial evaluation phase). Each binary choice was presented for 1.5 seconds, followed by fixation in a jittered interval of 3.5 seconds in average (SD = 2.05 seconds, range of 1.5 - 11.5 seconds, 0.5 seconds interval). A green square around the chosen item indicated the participant’s response. If the participant did not respond on time, the message “You must respond faster!” appeared for 0.5 second, followed by an interval fixation for the rest of the ITI duration. Each run started with 2 seconds of pre-run fixation and ended with a 6 second post-run fixation completion to the last run fixation.
6. ***Resting-state scan*** (8 minutes; inside the scanner.): Participants were instructed to rest with their eyes open while viewing a black screen. Two scans were performed: anterior posterior (AP) and posterior anterior (PA) acquisitions.
7. ***Memory task* (5 minutes; outside the scanner.):** 24 snacks from the probe task (12 Go and 12 NoGo items) and 24 ‘new’ snacks (which did not appear in the experiment until now and where chosen from the snacks dataset) were presented on the screen one by one for 3 seconds. Participants rated each item, on a 5-point confidence scale, whether it had appeared in the experiment and if it was associated with a cue. This part included 96 trials (24 old and 24 new snacks, 48 trials for each question). A green square around the chosen answer indicated participants’ responses. If the participant did not respond on time, the message “You must respond faster!” appeared for 0.5 second.
8. ***Repeated WTP task:* (15 minutes; outside the scanner.):** The same auction as step 1 was repeated outside the scanner to test for preference changes.
9. ***Eating questionnaire* (5 minutes; outside the scanner.):** At the end of the experiment, participants completed an adult eating behavior questionnaire (AEBQ; Hunot et al., 2016).

### Follow-up sessions (sessions 2 to 5)

In the follow-up session participants did not undergo training and were scanned with the following protocol: a. passive viewing task (see session 1, subsection 2 above) b. probe task (see session 1, subsection 5 above), c. resting-state (see session 1, subsection 6 above) and d. anatomical scans – MPRAGE and FLAIR. Outside the scanner, the participants performed the e. memory task (see session 1, subsection 7 above) and f. the WTP task (see session 1, subsection 1 above).

#### Cues

In the task 30% of the stimuli in the training task were coupled with a visual cue, displayed in the middle of the screen (see Training task, under Cue-approach training section). Participants were asked to press a button as fast as they could in response to the cue. The neutral cue consisted of two asterisks (see Figure 1C). The incentive-cue was a number in range 21 to 24, indicating a future winning prize. Participants in the incentive-cue group were informed that one trial would be selected randomly at the end of the experiment and that they would win the amount shown as a bonus (21-24 NIS, equal to 5.7-6.5 $).

### Statistical Analyses

To examine how difference in behavioral changes procedures affect functional connectivity, and to assess the long-term maintenance of these effects. Our analyses comprised four components: (1) behavioral analyses, to examine the task-related and longitudinal effects of different procedures on preference changes; (2) training-related analyses to explore how training (session 1) influenced subsequent functional connectivity (FC) and brain network architecture, as measured by the resting-state scan conducted immediately after training; (3) longitudinal-maintenance analysis to assess how the long-term effects of behavioral changes are reflected in functional connectivity; and (4) a functional connectivity marker for the preferences-changes parameter, both immediately after training and longitudinally.

#### Exclusion Criteria

Based on previous studies and as preregistered, we had several exclusion criteria. A total of 144 participants took part in this study. Of these participants, 37 were excluded from the analysis due to one of the following:

1. Disengagement during the training task:

- Two participants were excluded due to false alarms greater than 5%. A false alarm was defined as responding to a NoGo item during the training phase.
- Three participants were excluded due to missed trials greater than 10%. A miss trial was defined as a Go trial when the participant did not respond at all (1.5 seconds after image onset).
- Four participants were excluded due to the cue onset time dropping below 200ms at any time during the training phase. Whenever the participant failed to respond during the one-second image onset of a Go trial, the cue onset was lowered by 50ms (starting at 750ms). Each successful Go trial increased the onset of the cue by 16.67ms. Reaching a cue onset below 200ms indicated that the participant did not respond in many successive trials.
2. Technical issues:

- Four participants pressed the wrong button during the training or probe phases.
- Five participants were not scanned (or stopped in the middle of the scan) due to technical issues with the magnet.
- Ten participants stopped the scan in the middle.
- One participant was excluded due to neural findings.
- Eight participants were excluded due to other technical issues.

#### Long term behavioral change effect

For the behavioral analyses, we used the probe phase, whereby participants were asked for binary choices between Go and NoGo items with the similar initial willingness to pay. We tested whether the proportions of Go items chosen during the probe task exceeded random choice levels (50% of choosing Go items, odds ratio = 1), for each time point and group separately, by applying a logistic regression with choices as dependent variable and participants as random effect, following the approach used in previous CAT studies (Botvinik-Nezer et al., 2021; Salomon et al., 2020). We pooled together trials of high- and low-value items, as previous studies showed that the arbitrary separation of item-value is not a dominant feature in CAT (Botvinik-Nezer et al., 2021; Salomon et al., 2018)

To test the long-term effects of CAT, we used a repeated measures logistic regression model with choices as dependent variable, with group, session, and the interaction between group and session as independent variables. Participants were set as random effect for this analysis.

All statistical analyses were performed using the Python *statsmodels* and *scipy* packages (Seabold & Perktold, 2010) and R package *lmer4* (Bates et al., 2015).

#### Functional connectivity processing

In this paper we focused only on the resting-state data.

### Image Processing

Raw DICOM images were converted to NifTI format using dcm2nii toolbox. NifTI files were converted to BIDS format (Appelhoff et al., 2019).

To denoise the data, we first performed an independent component analysis (ICA) on each scan using the FSL Melodic toolbox, resulting in an unrestricted number of components. These components were classified as signal or noise using the FSL-FIX procedure (Griffanti et al., 2014; Salimi-Khorshidi et al., 2014). Components from twenty percent of the data (equally distributed between groups) was manually labeled as signal or noise and used as a training set for the FIX algorithm. Then, after the algorithm trained, the remaining images (80%) were automatically labeled. Images were denoised based on that classification and then registered to MNI 2mm template.

### Image parcellation

Images were parceled into 400 cortical regions using the 7-networks Schaefer’s atlas (Schaefer et al., 2018) and 16 sub-cortical regions using Pauli’s atlas (Pauli et al., 2018). Four cortical regions were excluded from further analysis due to low signal, based on low signal (Chopra et al., 2021). For each participant in each session (both runs combined), we generated a functional connectivity (FC) matrix by computing pairwise Pearson correlations between functional time-series extracted from those 412 brain regions using the Nilearn package for Python (Abraham et al., 2014), resulting in a 412×412 FC matrix for each scan, for each subject..

### Network-based statistic toolbox

The network-based statistic (NBS, <u>(Zalesky et al. 2010)</u>) is a statistical approach for connectome-wide inference. First, a statistical test is performed on each edge of the connectivity matrix. Edges exceeding a predefined threshold form connected subnetworks, whose sizes (number of edges implicates) are recorded. A permutation-based null distribution is then generated by randomly shuffling group labels and recalculating the size of the maximal subnetwork. Finally, the sizes of the observed subnetworks are compared to this distribution, and significance is determined based on the observed network size exceeding 95% of the nulls (ie. p <.05), effectively controlling for multiple comparisons using family-wise error correction at the level of networks connected regions. We used the NBS to test for three different effects: (1) differences between groups after training session; (2) differences between groups for long-term behavioral effect maintenance; and (3) FC-marker for behavioral effect.

#### Training-related FC analyses

Here we aimed to examine differences in FC between groups immediately after training, i.e. in the first meeting. We used the NBS (Zalesky et al., 2010) to test for statistical differences between groups, using t-test as the statistical test, with a primary component-forming threshold of *t* = 2.75, 1,000 permutations to create the null-distribution, and component-level significance of p<0.05, FWE corrected.

Additionally, for each participant’s FC matrix, we calculated several graph metrics that capture network architecture: (1) the eigenvector centrality of each brain region (Das, 2004) which quantifies the importance of a region within a network based on its connections with other regions, and the connections of those neighbours; and (2) the BOLD signal variability for each region which reflects the synchronization of the network and degree of coordinated activity between different brain regions (Wehrheim et al., 2024). These metrics were computed using *Networkx* package for Python (Hagberg et al., 2008), and Kolmogorov-Smirnov test were applied to examine statistical differences in the regional distributions of these metrics.

#### Maintenance-related FC analyses

To investigate how the long-term maintenance of preference changes was reflected in FC, we compared groups across time to assess the impact of procedure type (NC vs. IC) on FC. To examine differences in FC between groups as related to time-after-training, we applied the NBS to FC matrices, with FDR correction, 10,000 permutations, and tested interaction between time and group as regressor.

#### FC-marker for preference changes

We aimed to identify FC markers associated with behavioral changes, reflected in the proportion of Go items chosen during the probe phase. To investigate task-related behavioral markers, we used the NBS toolbox, using F-test as the statistical test, with f = 4.5, 1,000 permutations to create the null-distribution, and significance level of p<0.05, FEW corrected, to assess FC associated with probe scores, for each group separately. We examined which edges were shared between the two groups and which were distinct to each. For long-term analysis, we applied the NBS toolbox, using F-test as the statistical test, with f = 15, 1,000 permutations to create the null-distribution, and a component-wide significance level of p<0.01, corrected, to evaluate the combined effect of time-in-days and probe score for each group separately. We adjusted the statistical thresholds to balance sensitivity and interpretability, ensuring that the identified subnetworks were meaningful rather than an overwhelmingly large set of correlated edges.

### Data availability

The project had preregistered and can be found at: (https://osf.io/ag3ws/?view_only=9137520c951e4edba0aeee680d75c8b9)

## Results

### Behavioral results

We first examined whether participants preferred Go items over NoGo items, considering group, session, and their interactions. In the first session, similarly to previous CAT studies (Botvinik-Nezer et al., 2020, 2021; Salomon et al., 2018, 2020; Schonberg et al., 2014), participants showed enhanced preferences for Go items in both groups (neutral-cue: OR = 1.25, 95% CI = [1.08, 1.46], z = 2.984, p = 0.001; Incentive-cue: OR = 1.20, 95% CI = [1.04, 1.38], z = 2.525, p = 0.006), but with no significant differences between the groups (p-value = 0.64).

In the second meeting (one month after the first one), we observed maintenance for the neutral-cue (NC) group (OR = 1.24, 95% CI = [1.05, 1.46], z = 2.568, p = 0.005). No effect was detected for the incentive-cue (IC) group (OR = 0.97, 95% CI = [0.83, 1.14], z = −0.333, p = 0.370). Furthermore, we found significant differences between the groups (p = 0.035) in this session.

Three months after the first session (session number three), we found an elevation in preferences for Go items for the NC group (OR = 1.26, 95% CI = [1.04, 1.54], z = 2.388, p = 0.008) but not for IC group (OR = 1.04, 95% CI = [0.90, 1.21], z = 0.580, p = 0.281), but with no significant differences between the groups (p = 0.14). These follow-up results are consistent with previous studies (Botvinik-Nezer et al., 2020; Salomon et al., 2018; Schonberg et al., 2014).

The same trend was maintained at the fourth meeting (NC group: OR = 1.26, 95% CI = [1.03, 1.54], z = 2.369, p = 0.009; IC group: OR = 1.02, 95% CI = [0.83, 1.26], z = 0.243, p = 0.404) and fifth meeting (NC group: OR = 1.35, 95% CI = [1.11, 1.65], z = 3.178, p = 0.001; IC group: OR = 1.01, 95% CI = [0.81, 1.28], z = 0.134, p = 0.447). A trend in differences between the groups was observed for the last two sessions (4^th^ session: p = 0.075; 5^th^ session: p = 0.067) Repeated measure logistic regression revealed a significant decrease in proportion of preference changes from the first to the third meetings (p = 0.006). All results are shown in Figure 2.

**Figure 2:**
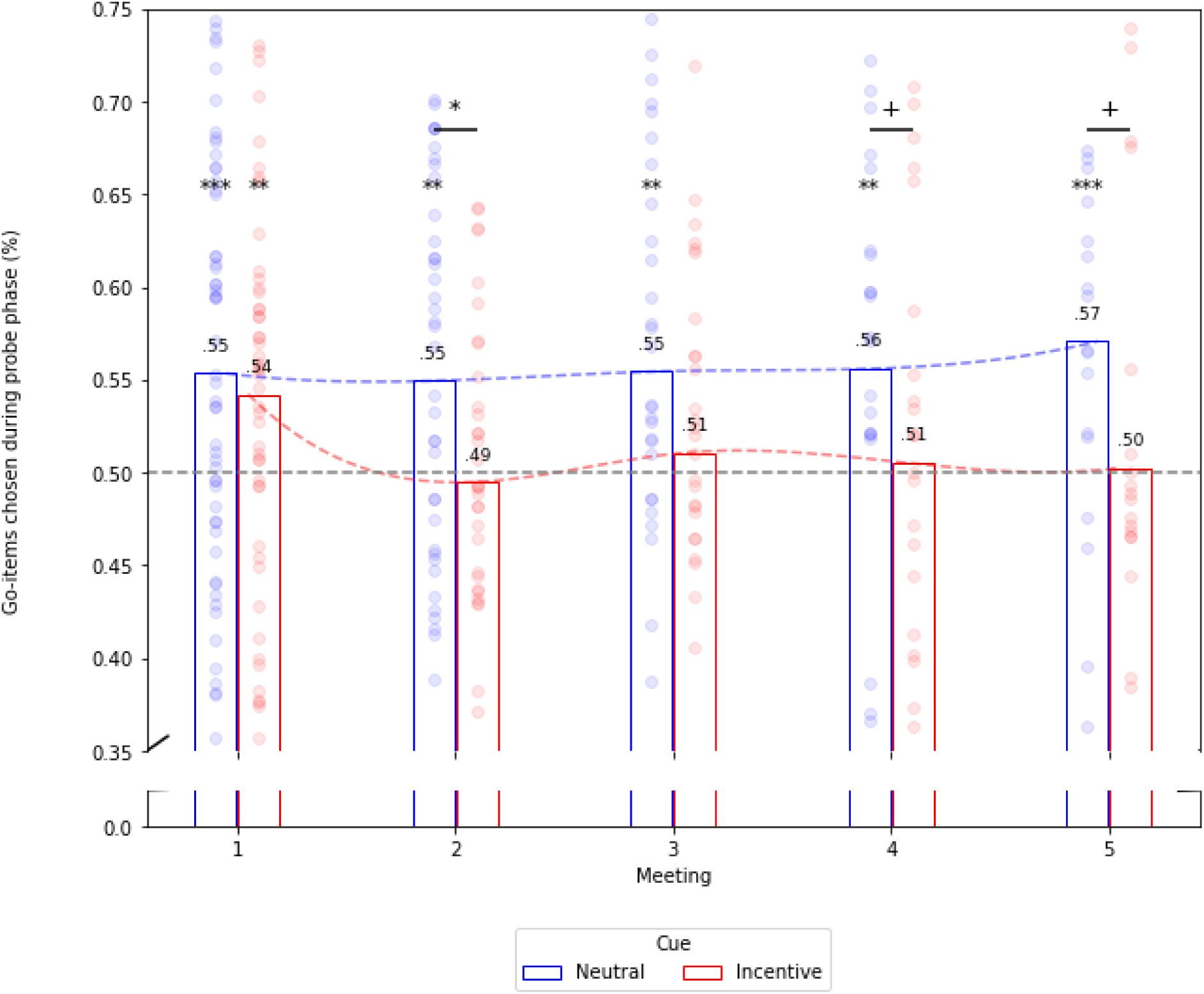
the proportion of Go items chosen (y-axis) separated to the different time points (x-axis). Blue bars refer to the neutral-cue group. The red bars refer to the Incentive-cue group. A gray dashed line indicates odds ratio of 1 (50%). Each dot represents a participant. Statistical significance is indicated p-value: < 0.1 = +; < 0.05 = *; < 0.01 = **; < 0.005 = ***. Numbers above the bars indicate the mean proportion of Go item chosen during probe for a given group in a given session.

### Training-Related Functional Connectivity

#### Statistical analysis for differences in FC

We initially examined differences in FC between groups for the first session, i.e. after the initial one-hour training session, to assess its impact under the neutral or incentive cue procedures. The NBS revealed a single significant component comprising three hundred and twenty six edges that connected 126 nodes (Figure 3a.). Out of these edges, 80 (24.5%) were connecting regions within the limbic network with itself, both raw and normalized data (Figure 3b.). A further 74 (22.7%) edges were assigned to be part of the DMN (see Figure 3a.b.c.). Next, we examined the FC differences between groups, by examining the mean FC at every significant edge. Most of the edges were showed higher FC in the IC group (96%, 312 edges) than the NC group. We found that in the NC group, network edges were primarily associated with the DMN and visual networks (see Figure 3c), while most connections involving limbic regions were classified under the IC group (see Figure 3b).

**Figure 3.**
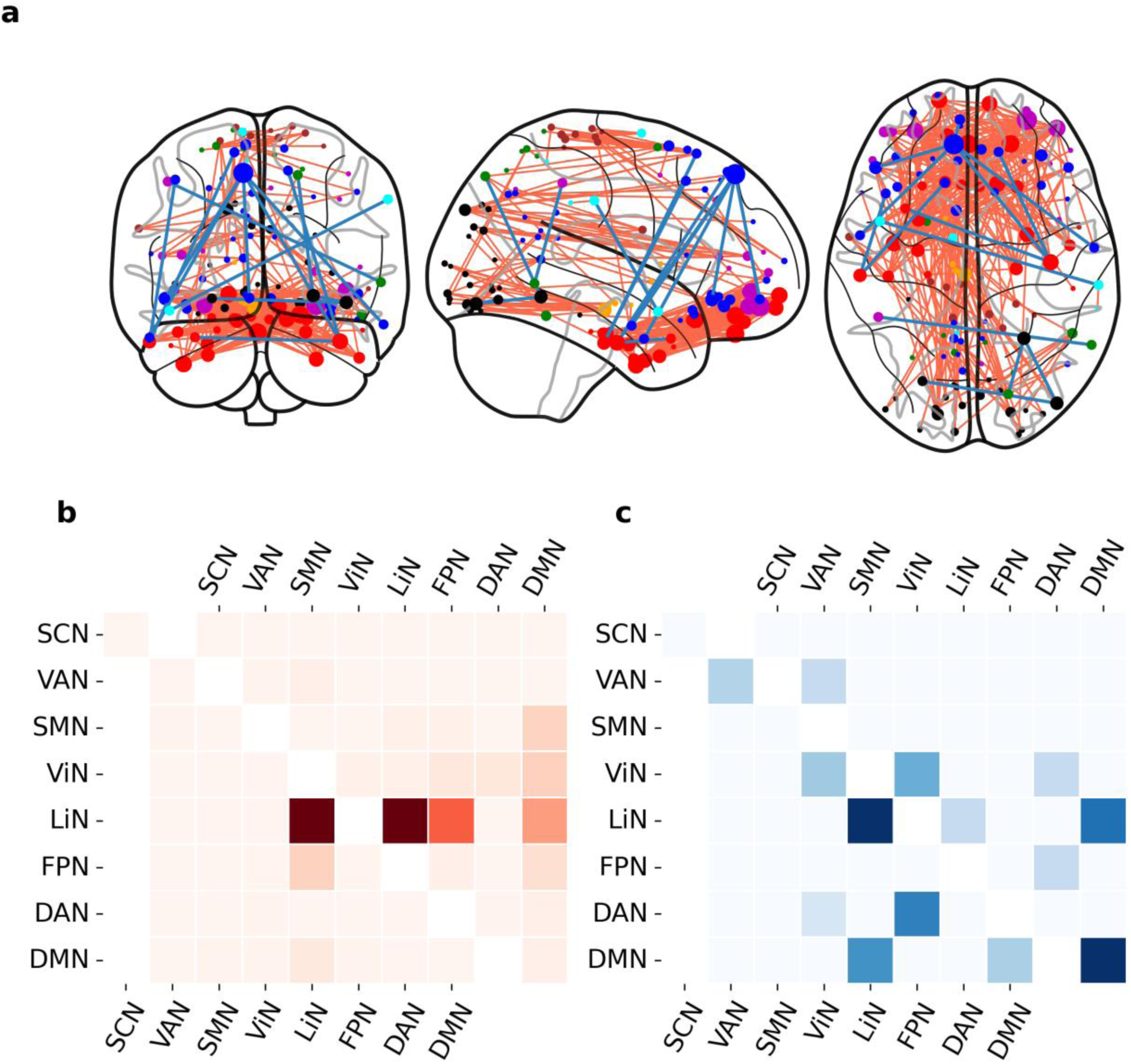
Visualization of network-based statistical results for differences in functional connectivity between neutral and incentive groups immediately after one-hour of training. Top row shows (a) connectome plots with node sizes reflecting the degree (number of significant edges), node colors representing network affiliations, and edge color representing group (blue – NC; red – IC). Middle row shows heatmaps of network-level connections. The upper triangle represents raw values, while the lower triangle represents normalized values across networks. Middle row left shows (b) Negative-intensity values, meaning higher FC for IC over NC. Middle row right shows (c) positive-intensity values, meaning higher FC for NC over IC. Network-coloring legend: subcortex – orange, Limbic – red, Frontoparietal network – purple, default mode network – blue, dorsal attention – green, ventral attention – cyan, Somatomotor – brown, visual - black.

#### Network architecture

We aimed to detect group differences in network centrality and synchronization, by calculating eigenvector centrality and BOLD-signal variability calculations for each node. Then, we compared the distributions of region-level values for each measure across canonical resting-state networks (Yeo et al., 2011). We found that eigenvector distribution, reflecting centrality, showed less network-centrality for the DMN in the NC group (p = 0.001). For network-synchronization (manifesting as low BOLD-signal variability), except for attention-networks (dorsal and ventral), all networks had higher network synchronization for the NC group. Results are shown in Figure 4.

**Figure 4.**
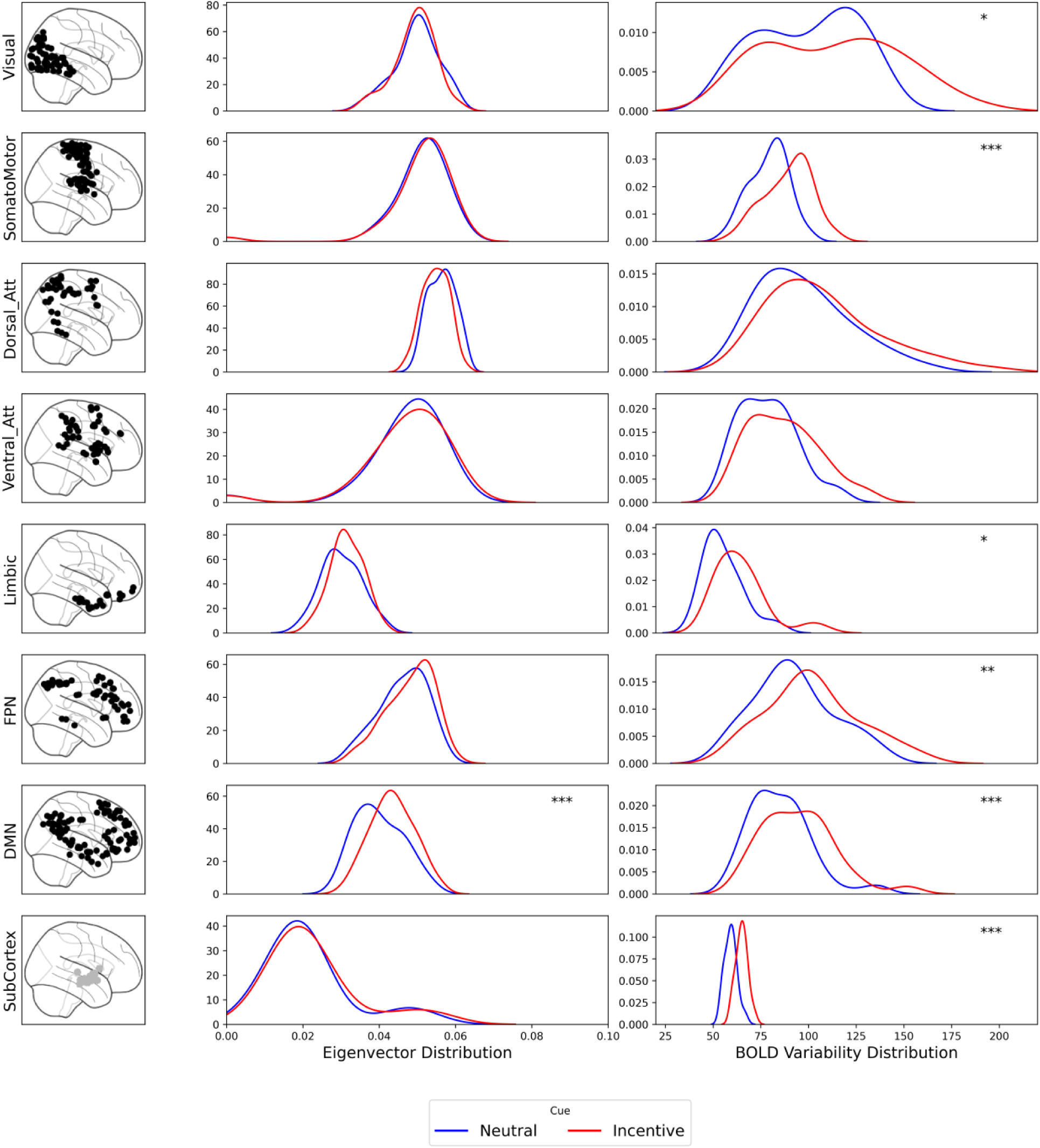
network architectures across groups and canonical networks immediately after one-hour training task. The figure showed the distribution of eigenvector as a centrality measurement (middle columns) and BOLD signal variability (right column) across different brain networks (left column) for the neutral-cue (blue) and incentive-cue (red) group. Each row corresponds to a specific functional network (from top to bottom): Visual, somatomotor, dorsal attention, ventral attention, Limbic, Frontoparietal, Default Mode Network, and subcortex. Asterisks denote statistical significance levels for group differences: + (p < 0.1), * (p < 0.05), ** (p < 0.01), *** (p < 0.005).

### Maintenance-related FC analyses

We next examined differences in FC between groups over time by testing the interaction between group (NC vs. IC) and time (timepoint 1-5). This analysis revealed a single statistically significant component comprising 91 significant edges connecting 88 regions. Out of these edges, 44 (48.4%) had higher FC in the NC compared to IC group. The mammillary nucleus was found to be a hub in both groups (high number of edges connected to this regions). We found that, for the NC group, most of the sub-cortex edges were connected with the somatomotor and dorsal attention networks. However, for the IC group, sub-cortex edges were mainly connected with limbic network regions. Finally, we observed that Frontoparietal regions predominantly associated with edges showing higher FC in the IC group, whereas DMN regions were more frequently linked to edges showing higher FC in the NC group. Results are shown in Figure 5.

**Figure 5.**
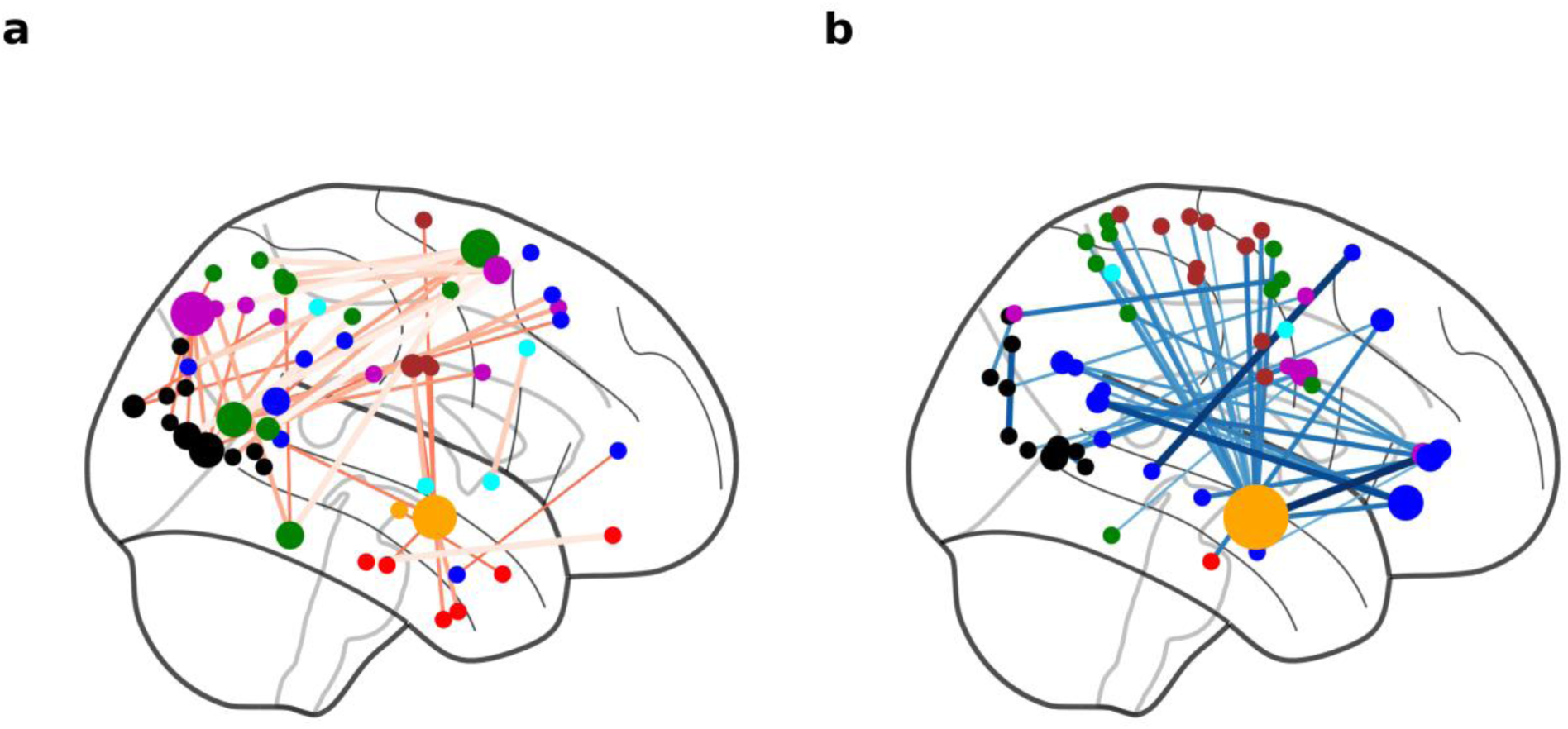
Modeling the interaction between group and session (in days). Node sizes reflecting the degree (number of significant edges) and node colors representing network affiliations. Network-coloring legend: subcortex – orange, Limbic – red, Frontoparietal network – purple, default mode network – blue, dorsal attention – green, ventral attention – cyan, Somatomotor – brown, visual - black. a) edges with higher functional connectivity for Incentive-cue groups. b) edges with higher functional connectivity for neutral-cue groups.

#### FC-marker for preference changes

Our next analysis examined resting-state FC correlates of behavioral changes, as indexed by the proportion of Go items chosen during probe phase of every testing session.

In the NC group, we identified a significant component comprising 39 edges connecting 49 regions that correlated with the behavioral parameter. In the IC group, we identified a component comprising 51 significant edges connecting 59 regions. Among these, 19 edges were common to both groups. The hypothalamus (HTH) emerged as a central hub in both the common and unique connectivity patterns. In the IC group, we observed a centralized network, where most edges connected the hypothalamus to the dorsal attention and Frontoparietal networks (see Figure 6a). In contrast, the unique connectivity pattern in the NC group was more distributed across multiple canonical networks (see Figure 6b). These results are consistent with the training-related resting-state FC analysis, showing less centrality for the NC group. Within the common pattern, the hypothalamus was linked to both ventral and dorsal attention networks. Also, for the common pattern, connections between visual and somatomotor regions were observed (see Figure 6c).

**Figure 6.**
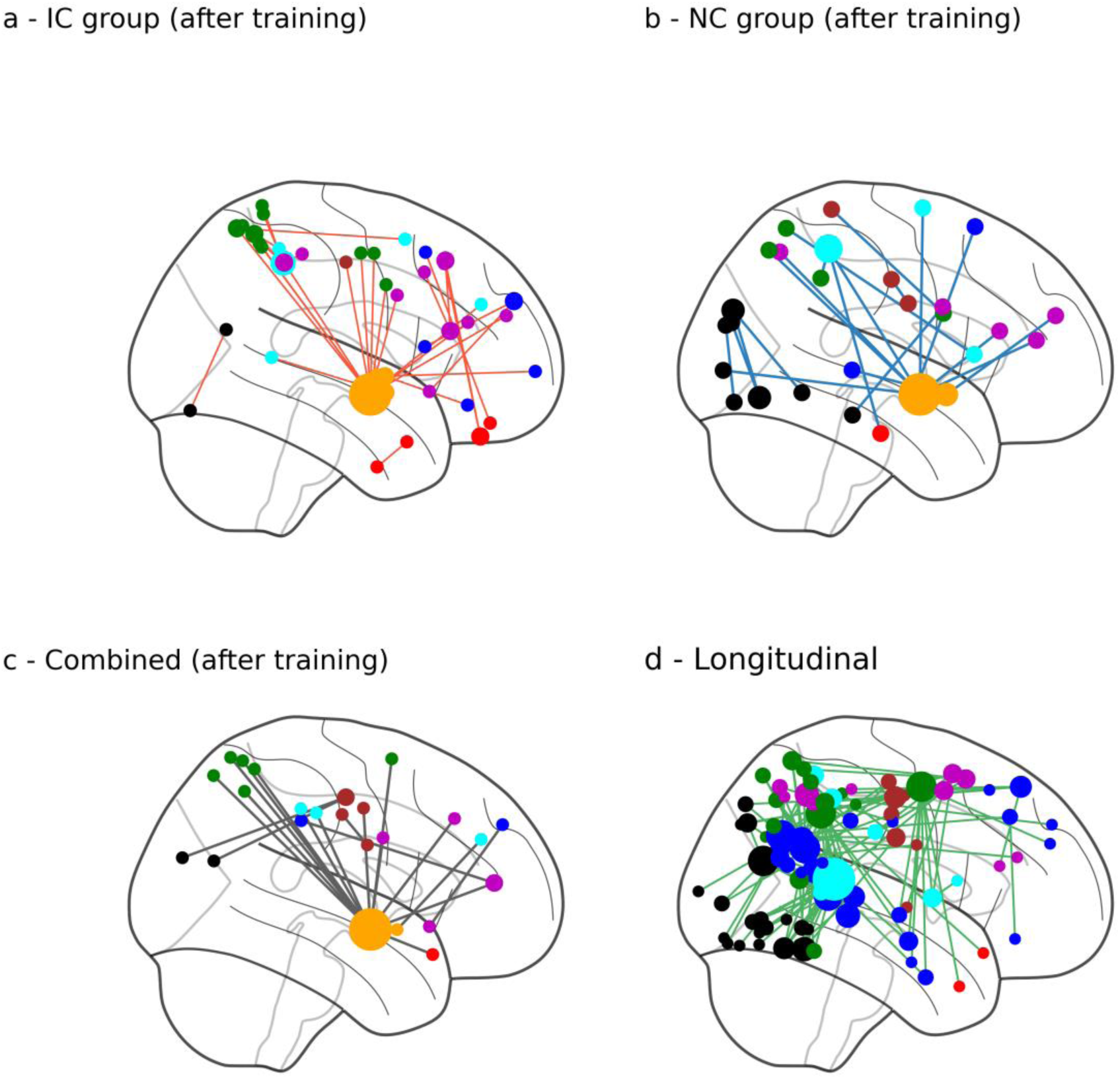
Functional-connectivity marker for behavioral changes. Node sizes reflecting the degree (number of significant edges) and node colors representing network affiliations. Network-coloring legend: subcortex – orange, Limbic – red, Frontoparietal network – purple, default mode network – blue, dorsal attention – green, ventral attention – cyan, Somatomotor – brown, visual - black. a) unique edges assigned to incentive-cue group after training session. b) unique edges assigned to neutral-cue group after training session. c) common edges for both incentive- and neutral-cue groups after one-hour training session. d) FC-marker for neutral-cue group for the combined effect of time and behavioral effect.

Next, we examined the maintenance-related FC marker for each group separately. In the IC group, no significant edges were found to correlate with time and behavioral effects, consistent with the behavioral results. In contrast, the NC group exhibited a significant component comprising 115 significant edges connecting 90 regions that was associated with the combined effect of time-in-days and behavioral changes. The temporal-occipital-parietal junction emerged as the main hub (16 connections, 14% of all significant edges). Strong connectivity was also observed in the DMN, dorsal attention, and visual networks (see Figure 6d).

## Discussion

In the current study, we used a novel design to directly examine neural differences between behavioral changes paradigms-non-external reinforcement preference change versus incentives. We used the cue-approach training (CAT, originated in (Schonberg et al., 2014)), a procedure for preference changes using the mere association of a speeded motor press response to a neutral cue. Two groups were tested: the “regular” non-reinforced neutral-cue group (NC), and the incentive-cue (IC) group in which the cue represented a monetary reward that participants will receive at the end of the experiment. Participants were scanned five times over one year. At the first assessment, the full CAT experiment was performed, included initial willingness to pay, training and probe. In follow-up sessions, no training was performed (meaning, only testing for preservation of behavioral effect). We tested both behavioral effects and functional connectivity (FC) differences between groups.

Behaviorally, both paradigms affected preferences immediately after training, as reflected by a higher proportion of chosen trained items over untrained items in the binary choice probe phase. Our hypothesis that the incentive-cue group would yield similar preference change rates compared to the neutral-cue group was confirmed. More interestingly, as we proposed in our preregistration (https://osf.io/ag3ws/?view_only=9137520c951e4edba0aeee680d75c8b9), the neutral-cue training maintained the preference change longer than the incentive training; that is, above-random choices were observed for neutral-cue group for each of the four follow-up meetings, an effect that was not observed in all follow-up meetings for the incentivized group. We propose that training during the CAT task internally reinforces action for specific items (Salomon et al., 2022), leading to higher selection rates during the probe phase. This putative internal reinforcement may have contributed to the long-term persistence of these preferences.

These behavioral results highlight the efficacy of non-external reinforcement paradigms in shaping preferences over extended periods, with implications for real-world behavioral changes. The sustained preference shifts observed in the neutral-cue group suggest that even in the absence of tangible rewards, simple motor associations can create lasting behavioral adaptations. Notably, while previous research has shown preference changes persisting for up to six months, our findings extend this duration to over a year, emphasizing the long-term impact of this mechanism. Understanding how non-externally reinforced behavior lead to persistent preferences could inform strategies in various real-life contexts.

Consequently, when examining differences in FC between groups immediately after the first and only training session conducted during the year, we found that most connections distinguishing the groups showed higher FC in the incentive-cue group, primarily within the limbic system. In contrast, the neutral-cue group exhibited stronger connections within the DMN. Graph analysis revealed a more centralized network architecture in the incentive-cue group, whereas the neutral-cue group displayed greater synchronization, reflected by lower BOLD signal variability. We propose that the differences in FC patterns between groups were driven by the type of cue. In the incentive-cue group, reinforcement was explicitly tied to monetary rewards, which activated reward and emotion-related regions (observed in the limbic system connections), directing the network along specific reward pathways (McClure et al., 2004;

Small et al., 2005; Weinstein, 2023). In contrast, the neutral-cue group may have relied on internal reinforcement at the participant level (Salomon et al., 2025). Without an external reward to guide their decisions, these participants were more likely to engage in flexible, self-directed reward-exploration. This exploratory behavior may have led to greater activation of the DMN, reflecting a more unbiased, internally driven process (Molnar-Szakacs & Uddin, 2013). The higher synchronization observed in the neutral-cue group may reflect an internally-driven reinforcement process, where decision-making relies more on distributed mechanisms, in contrast to specific reward-driven processes elicit by the incentive training. This hypothesis should be examined in a designated experiment.

When examining FC differences between groups over time, we found a sub-cortical region – the mammillary nucleus – to serve as a hub in the connectivity pattern in both groups. For the incentive-cue group, and correspondingly with the training-related FC pattern observed for this group, the limbic system was mainly connected to the mammillary nucleus. For the neutral-cue group, the same subcortical region connected mostly to the somatomotor, dorsal attention and DMN networks. The role and connectivity pattern of this reward- and memory-related mammillary body region (Coenen et al., 2018; Vann & Aggleton, 2004) may inform the underlying mechanism of preference changes maintenance. We suggest that for the incentive external reinforcement group, the brain putatively relied mostly on higher-level FC to maintain behavioral change, involving the limbic and the Frontoparietal networks. On the other hand, non-external reinforcement relies mostly on low-level FC, involving the somatomotor, dorsal (low-level) attention and the DMN. The observed result for neutral-cue group aligns with the Dorsal Value Pathway (DVP) hypothesis, and strengthens our suggestion of a pathway underlying non-reinforced behavioral changes (Schonberg & Katz, 2020).

Finally, when identifying an FC marker for behavioral changes, we found the hypothalamus functioned as a central hub for task-related FC changes. Recent findings suggest that the lateral hypothalamus plays a key role in shifting food-related behavior (Sharpe, 2024), which may explain its involvement in modifying snack-related preferences. When examining the maintenance of behavioral changes, a high number of connections were assigned to the DMN and the dorsal attention networks, indicating their role in sustaining behavioral adaptations over time. These results suggest that low-level connectivity patterns may contribute to the long-term maintenance of behavioral changes. Our findings provide evidence, albeit via reverse inference, for an internal reinforcement mechanism underlying the CAT paradigm, putatively driven by low-level network activation and the dorsal-value pathway hypothesis (Schonberg & Katz, 2020). The distinct functional connectivity patterns observed between the neutral- and incentive-cue groups suggest that different neural processes are at play. In the incentive training, the familiar monetary reward activates emotion and high-level regions, resulting in a more centralized network architecture. Conversely, the neutral training, which lacks a unified external reward, leads to a more distributed network, particularly in the DMN. This distributed architecture may reflect the brain’s search for individualized internal reinforcers. The long-term preservation of these effects, as seen in the follow-up sessions, further supports this hypothesis. The mammillary nuclei’s is a hub in both groups, but with distinct connectivity patterns, highlights the importance of memory processes in maintaining behavioral changes. For the IC group, connections primarily involved the limbic system, while the neutral group showed stronger connections between somatomotor, dorsal attention, and DMN networks. These patterns align with the proposed Dorsal Value Pathway, a proposed neural circuit underlying preference changes in the absence of external rewards (Schonberg & Katz, 2020). This internal reinforcement mechanism could explain why the neutral-cue training leads to more durable preference changes compared to external monetary rewards, as reflected in both behavioral outcomes and neural connectivity patterns.

The main limitation of our study is the drop-out rate of participants due to the COVID-19 pandemic, which in turn shrunk our sample size in the late follow-up sessions. We used mixed-effect models to control this issue, but our findings require replication in a larger sample. Future studies also could incorporate a pre-training resting-state scan to directly assess how different cue types during training influence functional connectivity, which an aspect not addressed in our design. Another limitation of our results concerning long term maintenance is that they could be explained by the “choice leads to choice” phenomenon, also known as choice-induced preference change, in which making a decision can alter individuals’ preferences, often making them more aligned with the choice they selected, creating a self-reinforcing cycle (Sharot et al., 2010). Moreover, this effect can persist for several years (Sharot et al., 2012). Thus, we cannot rule out the long term maintenance change was also driven by the initial choices. It should be noted that we have previously shown that the CAT-induced preference changes even when the first probe phase was performed three days after training (Botvinik-Nezer et al., 2021).

Our study sheds light on the longitudinal mechanism of preference changes, focusing mostly on how functional connectivity evolves over time, and offered an internal reinforcement mechanism to produce this behavioral effect. The internal reinforcement sustains longer than monetary (incentive) reward, in a way that reflected in either behavioral or neural connectome. Future studies can dive into changes in FC during training, enhancing and upscaling temporal and spatial resolutions.

To conclude, our study provides novel insights into the neural mechanisms underlying longitudinal preference changes, induced by external and internal reinforcers using modified version of the cue-approach training paradigm. While both neutral and incentive groups demonstrated significant behavioral effects after training, the neutral-cue group’s preferences were more durable over time. Functional connectivity analyses revealed distinct functional connectivity patterns between groups, highlighting the role of limbic and Frontoparietal networks in externally driven preference changes, contrasted by DMN, somatomotor and dorsal-attention networks associated with non-external reinforcement. These findings support the hypothesis of a dorsal-value pathway in non-external reinforcement learning and suggest that the internal reinforcement mechanism offers a basis for sustained preference changes over time. By advancing our understanding of reinforcement-based learning, this research may inform strategies to enhance long-term behavior modification in various applied settings.

## Acknowledgments

This work was supported by a grant from The Center for AI and Data Science at Tel Aviv University (TAD), a collaborative grant from Tel Aviv and Monash Universities (AF and TSc), and the European Research Council (ERC) under the European Union’s Horizon 2020 research and innovation programme (grant agreement n° 715016) granted to Tom Schonberg.

## Notes

### Competing Interest Statement

The authors have declared no competing interest.

